# Characterizing common loss-of-function genes and their potential utility in assessing population variability and chemical susceptibility

**DOI:** 10.64898/2025.12.16.694775

**Authors:** Chanhee Kim, Zhaohan Zhu, Abderrahmane Tagmount, W. Brad Barbazuk, Rhonda Bacher, Leah D Stuchal, Christopher J. Martyniuk, Christopher D. Vulpe

## Abstract

Inter-individual and population variability in susceptibility to chemical exposures confounds determination of threshold exposure levels to protect the most vulnerable. Current risk assessment frameworks, in the absence of empiric chemical-specific data, generally recommend default or probabilistic adjustment factors to account for such variability. We present an experimental approach to incorporate common genetic variants potentially impacting population-level differences in toxicant susceptibility into human cell-based models for any cellular apical endpoint of interest. We focus on the genes with the most common aggregate loss-of-function (LoF) alleles in the gnomAD v3.0 data which we designated as the PopVarLoF set. Unexpectedly, enrichment analysis of these genes found significant overrepresentation of gene products playing important functional roles in toxicology. Interrogation of GWAS and PheWAS databases found that these genes are associated with diverse metabolic phenotypes consistent with the relevance of the PopVarLoF set in studying variability of toxicant response in human populations. We further characterized the PopVarLoF set by developing custom lentiviral CRISPR knockout libraries targeting the PopVarLoF genes to assess their functional essentiality in the HepG2/C3A cell line. Functional disruption of 14 of the PopVarLoF genes (∼1 %) without toxicant exposure resulted in significant growth defects in this cell line, consistent with the majority of PopVarLoF gene products having non-essential roles. The development of human cell-based toxicity assays or other NAMs which include the empiric assessment of common genetic sources of population variability in susceptibility to chemical exposure could contribute to more robust risk assessment which protects vulnerable populations while reducing uncertainty.

**Impact statement:** We characterize common loss of function genetic variants which could impact toxicant susceptibility and describe an approach to incorporate them into NAMs to enable empiric estimates of the contribution of genetic variability to diverse toxicity endpoints.

## Introduction

Chemical risk can vary substantially from individual to individual and population to population. Inter-individual variation in response to toxicant exposure can be influenced by intrinsic and extrinsic factors including life stage, sex, inherited genetic variants, epigenetics, nutrition, as well as co- and cumulative exposures(Zeise et al., 2013; Birnbaum et al., 2016). Incorporating such variability between individuals and populations represents a considerable challenge in toxicological risk assessments(Miller et al., 2001; Rusyn et al., 2022; Lu et al., 2025; Petersen et al., 2025a). Of these multiple factors, the role of genetic variability in people to the risks associated with chemical exposures represents a particular risk assessment challenge. The combination of both direct genetic effects and gene-environment (GxE) interactions complicates the assessment of chemical risk, as individuals with certain genetic variants may experience adverse outcomes only under specific exposures or environmental conditions(Virolainen et al., 2023). Despite the recognized need—highlighted in recommendations from the National Academies of Sciences(National Research Council (US) Committee on Improving Risk Analysis Approaches Used by the U.S. EPA, 2009)—to incorporate population-level genetic variation into hazard identification and toxicity risk assessment, previous and ongoing efforts have been limited by our understanding of the genetic variation underlying susceptibility differences and methodologic challenges in implementing toxicity assays which consider genetic variation. In the absence of chemical specific empiric data, risk assessments have to rely on default adjustment factors or probabilistic models to address potential variability in susceptibility to toxicants(Anon, 2009; Petersen et al., 2025b). Targeted toxicity endpoint studies to assess the range of genetic susceptibility suggest that these approaches may over- or underestimate actual variability and thus risk(Varshavsky et al., 2023a). Integration of empiric estimates of genetic susceptibility to toxicants in NAMs could enable data-driven risk assessments and potentially reduce uncertainty(Lu et al., 2025) (Figure 1).

**Figure 1.**
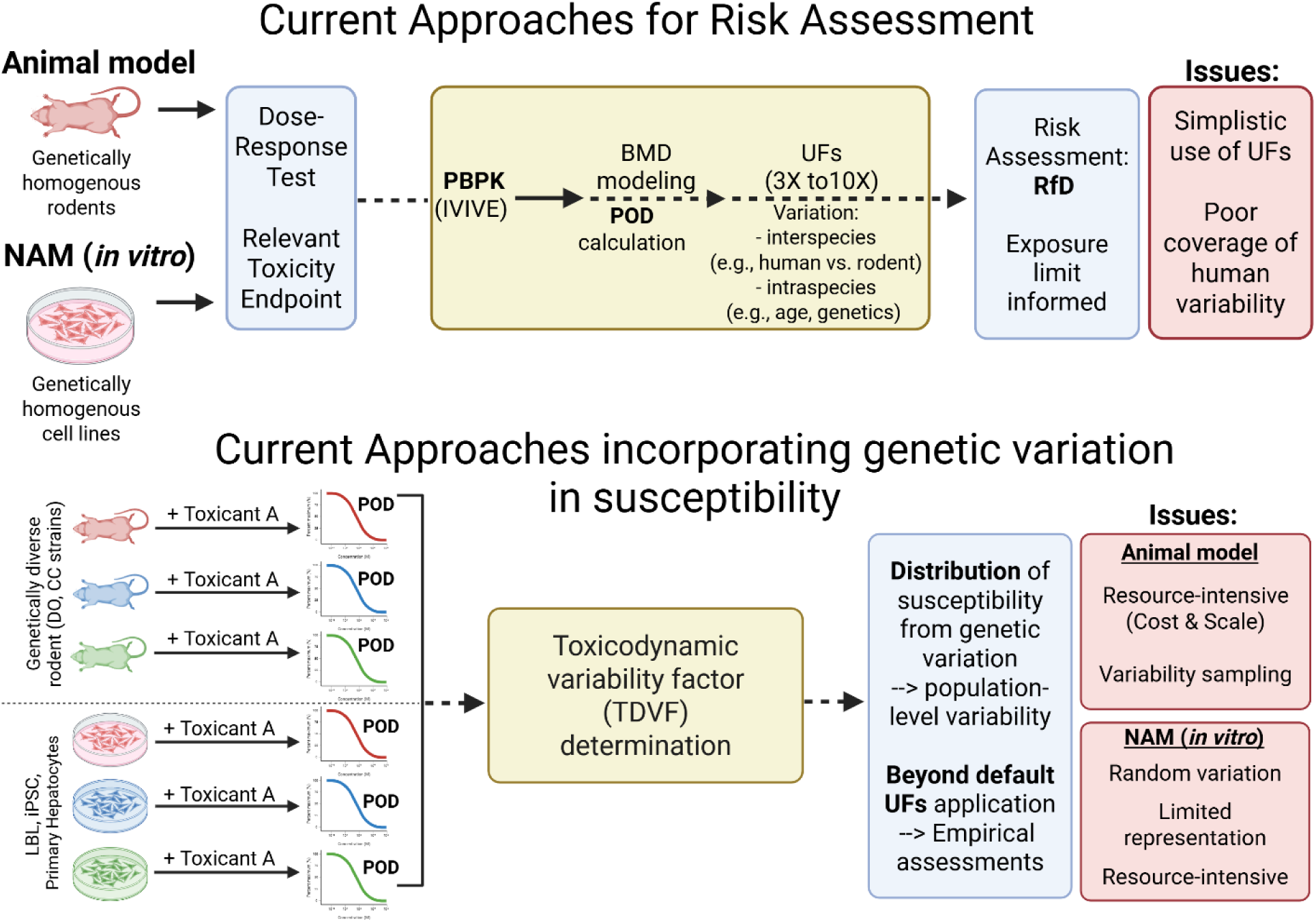
Current toxicological risk assessment framework and approaches to incorporate genetic variation in chemical susceptibility. NAM, new approach methodology; PBPK, Physiologically Based Pharmacokinetic; IVIVE, In Vitro to In Vivo Extrapolation; BMD, Benchmark dose; POD, point-of-departure; UFs, uncertainty factors; RfD, reference dose; DO, Diversity Outbred; CC, Collaborative Cross; LBL, Lymphoblastic lymphoma; iPSC, induced pluripotent stem cell.

Despite strong evidence that genetics play a role in toxicant susceptibility, our understanding of the specific genetic variants remains limited. This gap in knowledge, identified through studies in humans and model organisms, hinders our ability to incorporate genetic variability into risk assessments for toxic exposures(Anon, 2009). Targeted analysis of candidate genes previously identified to play a role in toxicology, such as ADME (absorption, distribution, metabolism, excretion), stress response, or cellular damage repair, have identified multiple genetic variants impacting toxicant response(Polonikov et al., 2009; Pinto and Dolan, 2011; Justenhoven, 2012; Zharinov et al., 2021; Carss et al., 2023). While providing important insights into the role of specific genes, these targeted approaches do not address the range of individual or population-level biological variation, and importantly, do not identify the most prevalent and functionally relevant variants driving differences in susceptibility. Similarly, multiple model organisms have contributed to our understanding of role of genetic variation in response to toxicants(Zhou et al., 2017; Dornbos and LaPres, 2018; Widmayer et al., 2022). Toxicant exposure studies in sets of inbred mice strains and recombinant inbred mice strains as well as more complex strain sets such as the Collaborative Cross (CC) populations and Diversity Outbred (DO) mice(Threadgill and Churchill, 2012) enabled efforts to sample and estimate variability in chemical metabolism, toxicokinetics, and toxicodynamics in mice (Threadgill et al., 2011; Churchill et al., 2012; Cichocki et al., 2017; Venkatratnam et al., 2017). However, while the overall population-level variation in response may be similar between mice and humans, the specific genetic variants driving these differences—particularly those responsible for outlier responses—often occur in different genes, produce distinct endpoints, and also presents the disadvantage of species-specific differences that can limit the direct translation of findings to humans(Harrill et al., 2009; Cichocki et al., 2017; van den Berg et al., 2021) (Figure 1). In people, to date, more systematic efforts to identify genetic variants that drive inter-individual differences in chemical susceptibility in human populations such as Genome Wide Association Studies (GWAS), and Phenome Wide Association Studies (PheWAS) have been hampered by the need for very large populations of individuals and complex environmental exposure histories (Ye et al., 2015a; Tam et al., 2019). As a result, while genetics is known to play a central role in individual and population-level risk, the field has struggled to identify the population-level relevant variants, with some notable exceptions, for most toxicants using genome-wide approaches(Motsinger-Reif et al., 2024).

To systematically address genetic variability in human populations and its potential role in differences in susceptibility to chemical exposure, our strategy is to take advantage of ongoing genome and exome sequencing efforts to identify common genetic variants in people and subsequently assess their impact on toxicant response. We contend that the most common predicted functional mutations in the human population(Karczewski et al., 2020a) represent a reasonable starting point, as these mutations could impact population level susceptibility. The Genome Aggregation Database (gnomAD)(Karczewski et al., 2020a) is currently the largest and most widely used publicly available collection of population genetic variation from genome and exome sequencing data(Gudmundsson et al., 2022). Computational prediction and aggregation of loss-of-function (pLoF) variants(Karczewski et al., 2020a; Minikel et al., 2020) into gene-level information previously identified a set of 1555 genes with an aggregate mean allele frequency (MAF) of pLoF variants of at least 0.1 % across all individuals in the database(Karczewski et al., 2020a). We contend that these genes represent a reasonable set to consider for a role in contributing to population-level variation in genetic susceptibility to toxicants.

Here, we characterize and experimentally evaluate the previously identified set of genes with an aggregate pLoF MAF of >0.1 % in gnomAD v3.0 which we designated as the PopVarLoF set (population variant loss of function gene set) and introduce a potential experimental framework for incorporating LOF variation into any human cell-based toxicity assays. We first annotated the PopVarLoF set using existing biological pathways databases. Then, the PopVarLoF set were interrogated for associations with human phenotypes in existing GWAS- and PheWAS-based analyses. Lastly, we generated a ‘synthetic’ population (thanks to Dr. Ivan Rusyn for this designation) of cells in which the PopVarLoF set of genes were individually disrupted, thus phenocopying homozygous LoF alleles, using CRISPR-based screening approaches to simultaneously assess their essentiality in a representative human cell line. We discuss a potential conceptual framework and experimental platform to incorporate common LoF variants into human cell-based assays to enable empiric estimates of genetic variability for diverse phenotypic endpoints used in high-throughput NAMs-based toxicological risk assessment.

## Materials and Methods

### 1. PopVarLoF genes

The Genome aggregation database (gnomAD v3.0) was previously used to identify 1555 genes with an aggregated mean allele frequency of predicted loss-of-function (pLoF) variants of greater than 0.1 % (Table S1) (Karczewski et al., 2020a). We removed misannotated and non-coding genes and designated the remaining 1551 genes as the PopVarLoF gene set for this study (Figure 2A).

**Figure 2.**
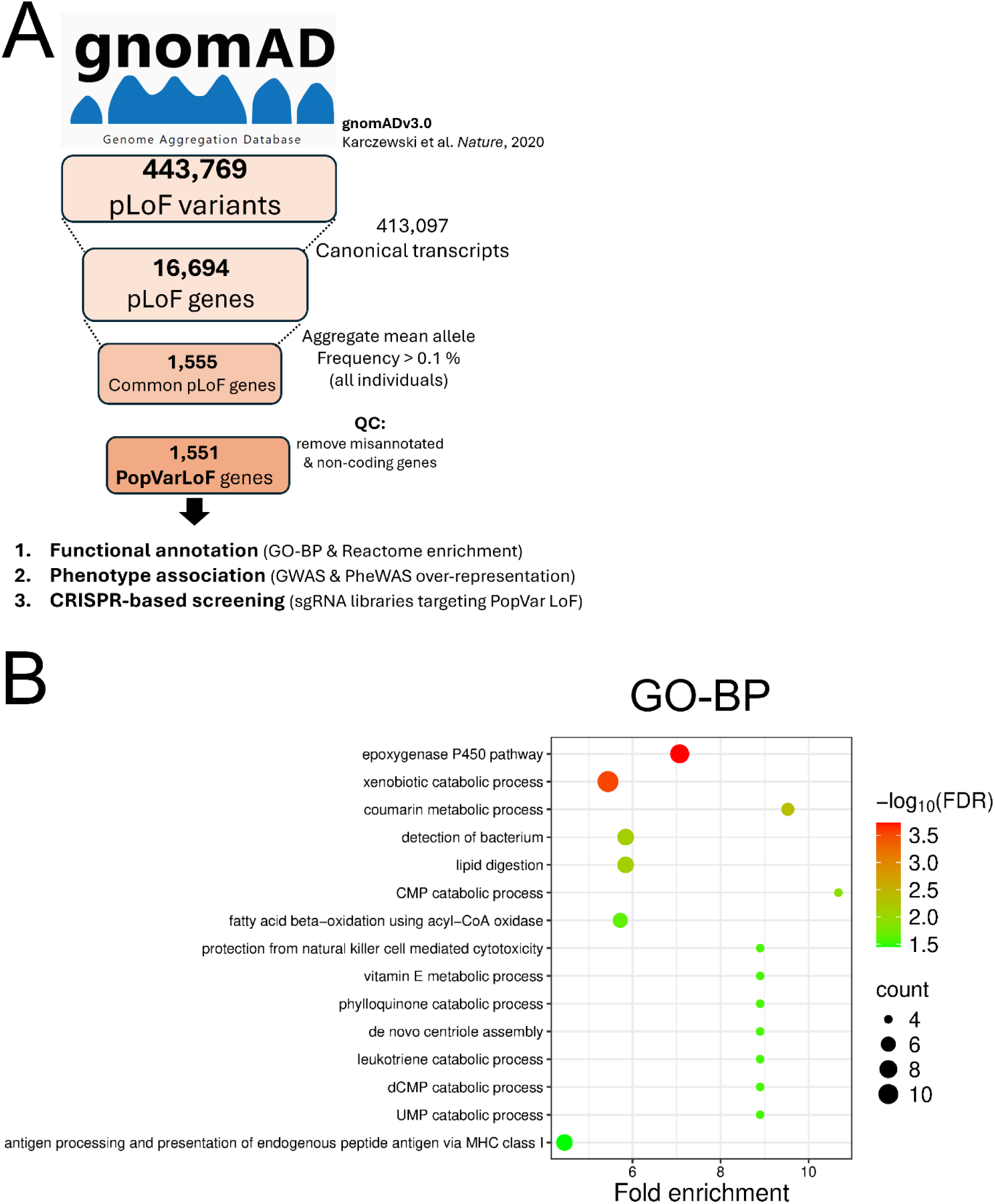
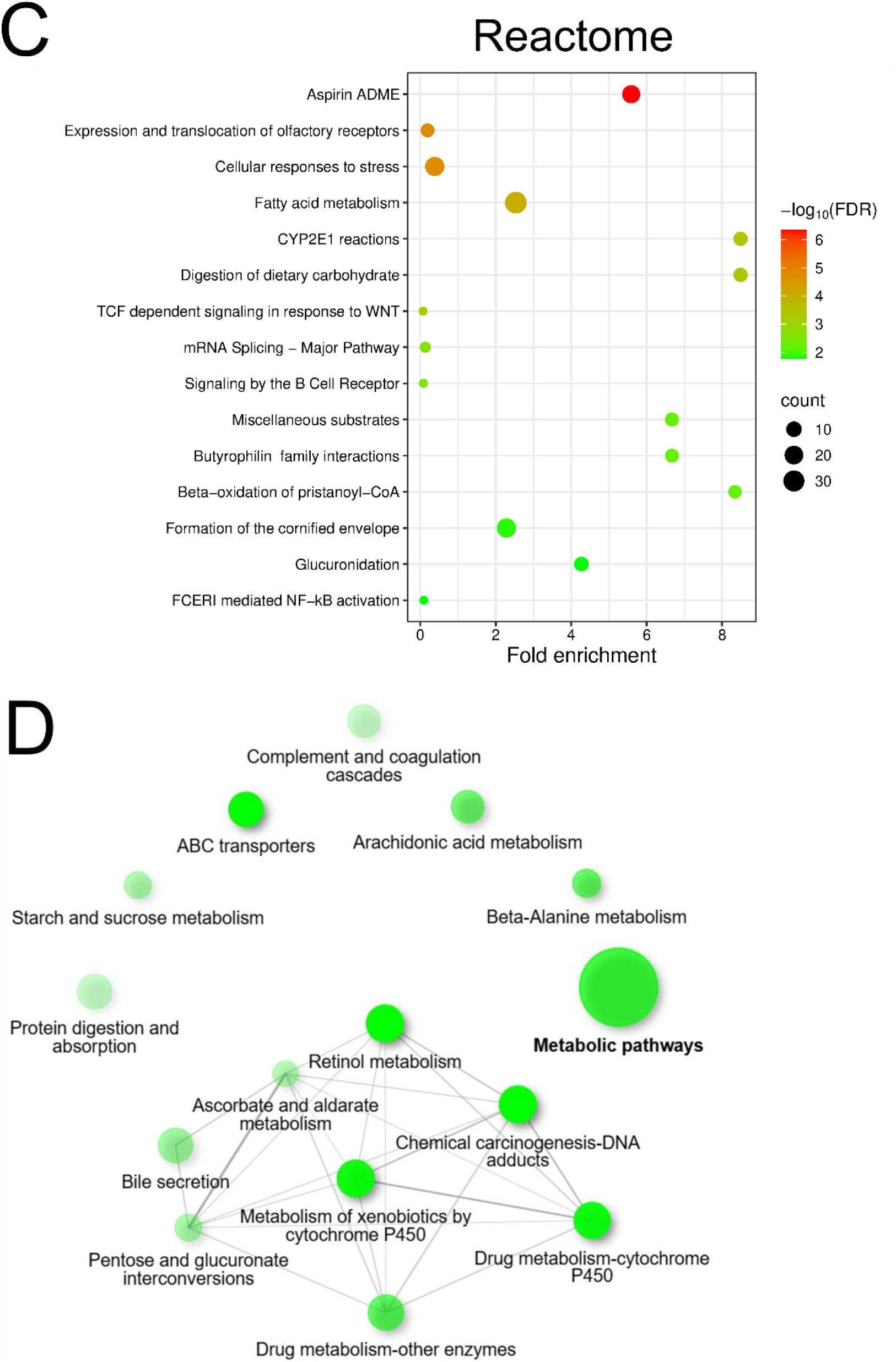
Identification and characterization of the PopVarLoF genes. (A) Overall schematic of PopVarLoF approach. Genes with aggregate mean allele frequency of >0.1 % of loss of function mutations were previously identified in gnomAD v3.0, filtered to remove non-coding genes, reannotated, and the resulting genes were designated as the PopVarLoF set. pLoF, predicted loss-of-function; QC, quality control; ED_50_, Median Effective Dose; (B) presents Gene Ontology (Biological Process, GO-BP) and (C) Reactome pathway enrichment of the PopVarLoF gene set, respectively, with top 15 terms being shown. The y-axis represents grouped terms of GO-BP and Reactome with x-axis being Fold enrichment and y-axis being corresponding terms. The circle size corresponds to number of genes in each category, and the color indicates false discovery rate (FDR) values as depicted in the right-hand side to the plot. (D) visualizes clusters of the most significantly enriched top 15 KEGG pathways.

### 2. Chromosomal location and functional enrichment analysis of the PopVarLoF gene set

The genomic distribution of individual PopVarLoF genes were visualized using Genome analysis tool provided by ShinyGO ([ShinyGO 0.82](http://bioinformatics.sdstate.edu/go/)). The PopVarLoF genes were then analyzed for functional enrichment annotation using Term Enrichment in AmiGO 2 ([AmiGO 2](https://amigo.geneontology.org/amigo)) which was subsequently conducted by PANTHER (pantherdb.org, accessed June 2025). We carried out enrichment analysis with GO-BP and Reactome databases (using default settings, accessed June 2025) and present the most specific terms significantly associated with the gene set using SRplot ([SRplot - Science and Research online plot](https://bioinformatics.com.cn/en)). KEGG pathway enrichment analysis was conducted via STRING (https://string-db.org/) and Cytoscape ([Cytoscape: An Open Source Platform for Complex Network Analysis and Visualization](https://cytoscape.org/)) for visualization of the enrichment.

### 3. GWAS Trait enrichment analysis

We used the GWAS Catalog (v1.0.2-associations_e113_r2024-11-03) and removed entries lacking mappable SNPs or gene annotations. Unmapped SNPs were assigned to the nearest gene using GRCh38 with the following R packages (SNPlocs.Hsapiens.dbSNP144.GRCh38, TxDb.Hsapiens.UCSC.hg38.knownGene, and org.Hs.eg.dbSee.) Traits were standardized using the “Mapped Trait” field in the GWAS catalog or if unavailable, the original disease trait was used. Studies reporting >10 traits, >1,000 genes, or traits with <5 genes were excluded. We analyzed enrichment of trait associations reported for the 1,469 of 1,551 genes in the PopVar set which matched HGNC names or aliases. For estimation of LoF variants in the 1469 genes as compared to random sets of genes, pLoF mutations were identified by Ensembl annotations (e.g., “stop_gained,” “frameshift_variant”). For the trait enrichment analysis, only intragenic pLoF SNPs were included (n=1,289 LoF, intergenic). Enrichment was assessed by comparing observed LoF gene-trait associations to 1,000 random intragenic gene sets (n=1,289). Only unique gene-trait pairs were counted to avoid redundancy. Trait-level z-scores were computed: z = (Observed PopVarLoF – Mean Random) / SD (Random), with two-sided p-values adjusted using FDR. Traits with ≥5 LoF hits and |z| > 2 were visualized.

### 4. PheWAS Trait enrichment analysis

All Data (UKBTOPMED, KoGES, MGIBioVU, and FinMetSeq), including PhenoStrings, were downloaded from https://pheweb.sph.umich.edu/ and https://koges.leelabsg.org/. If the broad category of SNP consequence was available, that data is shown as well. For each PheWAS dataset, we first identified phenotypes significantly associated with intragenic variants in the PopVarLoF gene set (for the 1551 annotated genes). We then sampled 1,000 random sets of associations between phenotypes and the PopVar genes for comparison by randomly selecting the same number of phenotypes from the whole dataset as the PopVarLoF gene set.

We estimated the proportion of a given phenotype category in each of the 1,000 random sets. We plotted the distribution of proportions and determined the mean and standard deviation (SD) for each phenotypic category. We calculated a z-score to assess the significance of the phenotypes associated with the PopVar gene set in PheWAS studies as compared to comparable random sets of phenotypes: Z = [Observed (PopVarLoF) – Mean (Random)] / SD (Random), with two-sided p-values adjusted using FDR. Traits with ≥5 LoF hits and |z| > 2 were visualized.

### 5. PopVarLoF sgRNA library preparation and quality check

We generated two sgRNA libraries targeting the PopVarLoF genes. 1555 genes with pLoF >0.1% were previously identified from gnomAD v3.0(Karczewski et al., 2020a). We assessed the current annotation at the time of initial study of each of these genes and removed 4 genes that were mis-annotated for the total gene set of 1551 genes. In order to identify sgRNA for a targeted CRISPR library, we selected the two top ranked sgRNA from a previous study(Gonçalves et al., 2021) that evaluated screening data for genome-wide CRISPR knockout (KO) libraries to rank the sgRNA for likely effectiveness. We noted that some sgRNAs were mis-annotated in the original genome wide libraries from which the sgRNA were identified and several genes in our set of genes of interest did not have sgRNAs identified. For these genes, we used a domain targeted sgRNA algorithm(Shi et al., 2015) to identify two sgRNA for each. We thus compiled two candidate sgRNA for each gene and as further quality control, we mapped each candidate sgRNA back to the human genome to confirm targeting location. The SET1 contains one sgRNA/gene (1551 genes targeted) with 100 non-targeting controls and 5 sgRNA targeting “safe harbor” gene(Aznauryan et al., 2022). Non-targeting sgRNAs were pulled from Minimal Genome Library(Gonçalves et al., 2021) while sgRNAs targeting safe harbor genes were two sites of Rogi1 and Rogi2 genes(Aznauryan et al., 2022). Likewise, the SET2 consists of sgRNAs targeting the same 1551 genes and controls as used for SET1, but at different locations. The SET1 and SET2 PopVarLoF sgRNA libraries were synthesized (Twist Biosciences) and the resulting oligo pools (SET1-oligo and SET2-oligo) were cloned into LentiCRISPRv2_Puro (Addgene #52961) separately as previously described(Shalem et al., 2014; Sobh, Loguinov, Yazici, et al., 2019) (See Table S1 for the complete list of sgRNA sequences). (Illumina NovaSeqX). We assessed the sgRNA representation before and after cloning (oligo pool vs. plasmid pool) by NGS of each pool, and demonstrated that the distribution of sgRNA was similar between the PopVarLoF-oligo (prior to cloning) and PopVarLoF -cloned (post-cloning) samples, with a Pearson correlation coefficient (r) of 0.9287 and 0.9008 for SET1 and SET2, respectively (Supplementary Figure S1A and S1B, respectively). We evaluated the histogram of sgRNA distribution and cumulative sgRNA read counts to assess representation. A six-fold difference between the 90^th^ and 10^th^ percentile has been suggested as the minimum acceptable distribution of sgRNA representation(Wang et al., 2014; Chen et al., 2019) and our libraries met this criteria (Supplementary Figure S1).

### 4. Time-course CRISPR functional essentiality screens

Lentivirus production was carried out as previously described(Sobh, Loguinov, Stornetta, et al., 2019). HEK293T cells were used to produce lentiviruses by co-transfection of either PopVarLoF SET1 or SET2 library plasmids, the packaging plasmid psPAX2 (#12260, Addgene), and envelope plasmid pMD2.G (#12259, Addgene). Both PopVarLoF SET1 and SET2 lentiviral libraries were functionally tittered in HepG2/C3A cells to determine the amount of virus required to obtain a multiplicity of infection of 0.3-0.5. For large-scale transduction for the screens, HepG2/C3A cells were seeded in four 12-well culture plates (1x10^6^ cells/well). After 24 h incubation, polybrene (Sigma) was added at a concentration of 8 µg/ml along with 0.4 µl (MOI= 0.3) of both the titrated PopVarLoF SET1 and SET2 lentiviral libraries in each well, followed by centrifugation (spinfection) at 33 ◦C at 1000xG for 2 h. After further incubation at 37 ◦C for 30 h, the lentiviral mix was replaced with 1 mL of the growth media in each well. After 48 h recovery period, the pooled cells were treated with puromycin (2 μg/mL) for 4 days to enrich transduced cells before time-course essentiality screens, resulting in almost no cells in the non-transduced control flask (initial KO population, T_0_). Consequently, HepG2/C3A mutant cells transduced with the complete PopVarLoF lentiviral library (SET1+SET2) were generated and cultivated for 14 days (7 doublings, T_7_) and 28 days (14 doublings, T_14_)(Russo et al., 2020; Zhao et al., 2021).

### 5. DNA extraction, library preparation, and next-generation sequencing

Genomic DNA of T_0_, T_7_, and T_14_ samples were extracted from 1.6x10^6^ cells (500X coverage) of each sample using the Quick-DNA Midiprep Plus Kit (ZYMO Research) following the manufacturer’s protocols. Amplicons for NGS Illumina sequencing were generated using the pairs of universal CRISPR-FOR1 forward primer and various CRISPR-REV**#** reverse primers (**#**: 1 to 48) specific for each sample (Table S2). Amplicons amplifying the sgRNA region in each sample were then pooled and gel purified using the QIAquick Gel Extraction Kit (Qiagen) and quantified using the Qubit HS dsDNA assay (Thermo Scientific). Equimolar amounts of each amplicon library were multiplexed in one pool. The NGS was carried out at the Interdisciplinary Center for Biotechnology Research (ICBR), University of Florida at Gainesville, using the NovaSeqX paired 150 bp high-throughput platform (Illumina).

### 6. Data processing and bioinformatics of time-course CRISPR screening data

#### 6.1. Data processing

All FASTQ files were first assessed for sequencing quality using FastQC, and summary reports were generated using MultiQC. Following the evaluation of sequencing quality, read counts were generated using the MaGeCK count command, which aligns reads to the PopVarLoF sgRNA library and produces a sgRNA-level count matrix used in downstream analysis. Additional quality control steps were applied to the read counts. We first evaluated sample-level read mapping efficiency and highly variable or low mapping percentages that may present a concern. Next, we check the count distribution across samples, and principal component analysis (PCA) to assess sample clustering and detect potential batch effects.

#### 6.2. Bioinformatics

Differential selection analysis was performed using DESeq2 (version 1.40.2) (Love et al., 2014) in R (version 4.3.3) to identify differentially selected sgRNAs between the baseline (T_0_) and time-course points (Day 14 and Day 28). For each comparison (Day 14 vs. T_0_ and Day 28 vs. T_0_), three T_0_ samples and four time-course samples were analyzed. The DESeq2 library size factor for normalization was calculated using the estimateSizeFactors function with the safe-harbor sgRNAs set as the control genes. Safe-harbor control sgRNAs have been demonstrated as a more appropriate baseline for normalization(Chen et al., 2018). Differential selection was assessed using DESeq2’s negative binomial generalized linear model with the Safe-harbor-based library sizes included as an offset. Adjusted p-values were calculated using the Benjamini–Hochberg procedure, and sgRNAs with an adjusted p-value < 0.05 were considered significantly differentially selected. A gene was deemed significant if at least one of its two sgRNAs was significant.

## Results

### 1. Genomic location and characterization of predicted LoF mutation in PopVarLoF set

We mapped the genomic localization and distribution of the PopVarLoF genes and did not identify any clear bias in chromosomal location (Figure S2). We analyzed the SNP data in GWAS catalog(Cerezo et al., 2025) and found over-representations of Stop-gained, Frame-shifted, Splice donor, and Splice acceptor variant categories in the PopVarLoF set as compared to other genes in the genome (Figure 3). Conversely, variant categories of Missense, Splice region, Splice donor region, Splice donor 5^th^ base were under-represented in the PopVarLoF genes. These findings of overrepresentation of pLoF variants in the PopVarLoF gene set in the GWAS SNP data is consistent with previous findings in gnomAD v3.0(Karczewski et al., 2020a).

**Figure 3.**
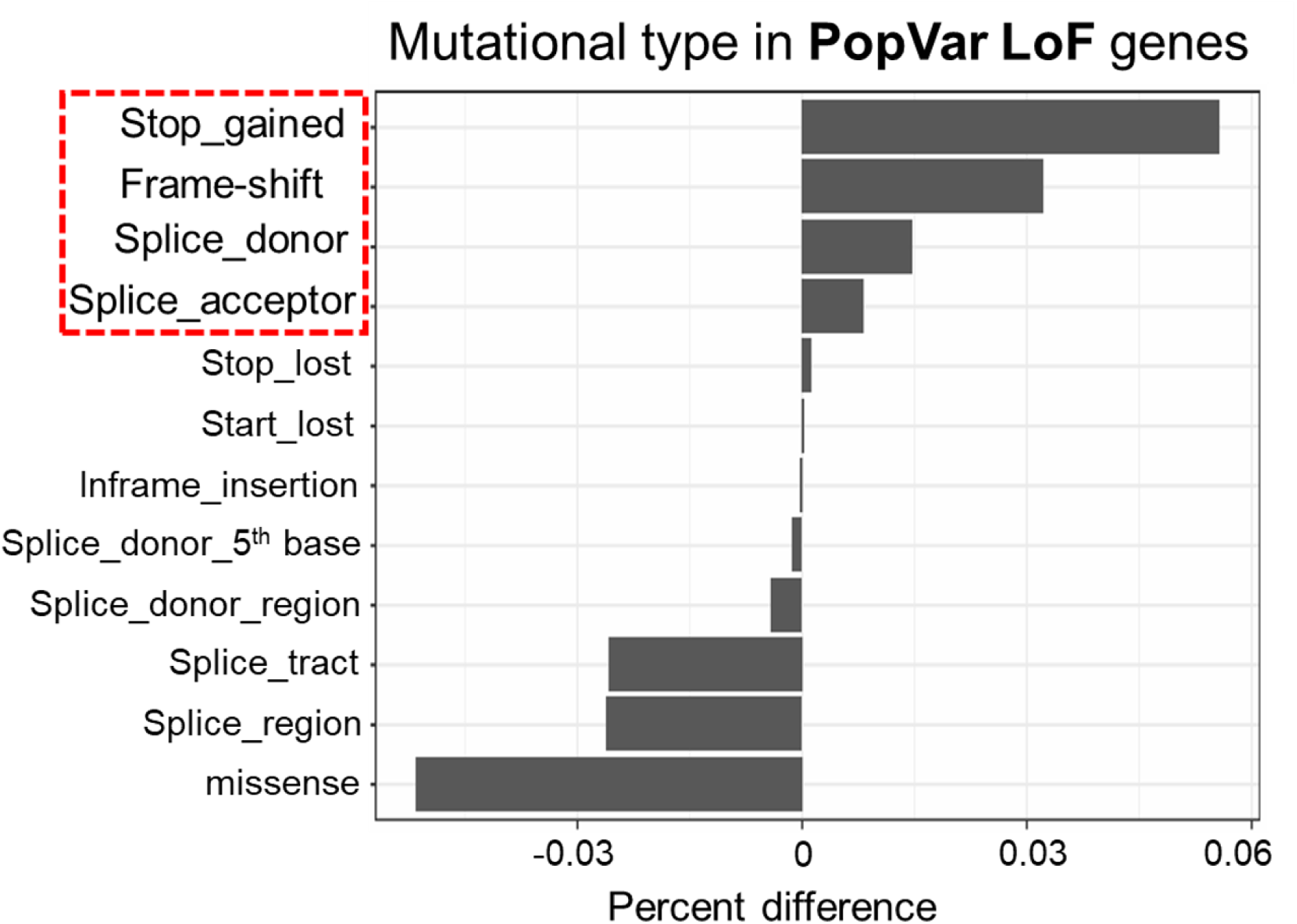
Frequency of mutational type in the PopVarLoF genes in the GWAS catalog. The x-axis indicates the percent difference of the frequency of each mutational type in the PopVarLoF set as compared to other genes in the GWAS catalog. Each mutational type is displayed in the y-axis.

### 2. PopVarLoF genes are enriched with metabolism and toxicology-related genes

We evaluated the PopVarLoF gene set for functional enrichment (the full list of 1551 PopVarLoF genes and their categorization for toxicological relevance are available in Table S1) including Gene ontology (Biological process; GO-BP) and Reactome. 11 of the top non-redundant 15 enriched GO-BP terms are related to metabolism, such as epoxygenase P450 pathway, xenobiotic catabolic pathway, coumarin metabolic process, and fatty acid beta-oxidation using acyl-CoA oxidase, others (Figure 2B, Table S5). GO-BP terms related to immunity and immune response were also enriched in the PopVarLoF genes, which included detection of bacterium, protection from natural killer cell mediated cytotoxicity, leukotriene catabolic process, and antigen processing and presentation of endogenous peptide antigen via MHC class I (Figure 2B, Table S5). Reactome analysis also indicated enrichment of our PopVarLoF genes in metabolism-related pathways such as Aspirin ADME, cellular response to stress, CYP2E1 reactions, Fatty acid metabolism, Beta-oxidation of pristanoyl-CoA, and others (Figure 2C, Table S5). We highlight the PopVarLoF genes encoding gene products involved in xenobiotic metabolism, transport and regulation in Table S5 and the immune related gene and gene products in Table S5. Furthermore, KEGG analysis showed a significant enrichment of metabolic pathways and other metabolism-related pathways, including ABC transporters, Retinol metabolism, drug/xenobiotic metabolism by CYP450 (Figure 2D). Together these findings indicate over-representation of genes involved in toxicity metabolism and immune function in the PopVarLoF gene set.

### 3. Over-representation of GWAS-linked metabolic phenotypes

We identified over-represented GWAS phenotypes associated with the PopVarLoF set of genes. The top 10 GWAS phenotypes (significant over-representation criteria: Z-score > 2 and p-adj. < 0.05, the entire 58 significantly associated traits are available in Table S6) associated with the PopVarLoF gene are shown in Figure 4. Seven of the top 10 over-represented GWAS phenotypes, including intrahepatic cholestasis of pregnancy (Z-score = 9.1), serum metabolite measurement (Z-score = 9.0), eicosanoids measurement (Z-score = 8.2), metabolite measurement (Z-score = 7.9), urinary metabolite measurement (Z-score = 7.7), bilirubin measurement (Z-score = 6.0), response to triamcinolone acetonide (Z-score = 5.8), and response to anticonvulsant (Z-score = 5.8) were related to metabolism (Figure 4A). We compared the individual PopVarLoF set of genes responsible for each GWAS association outcomes and found that distinct genes from the PopVarLoF set were responsible for each GWAS association with each trait (Figure 4B).

**Figure 4.**
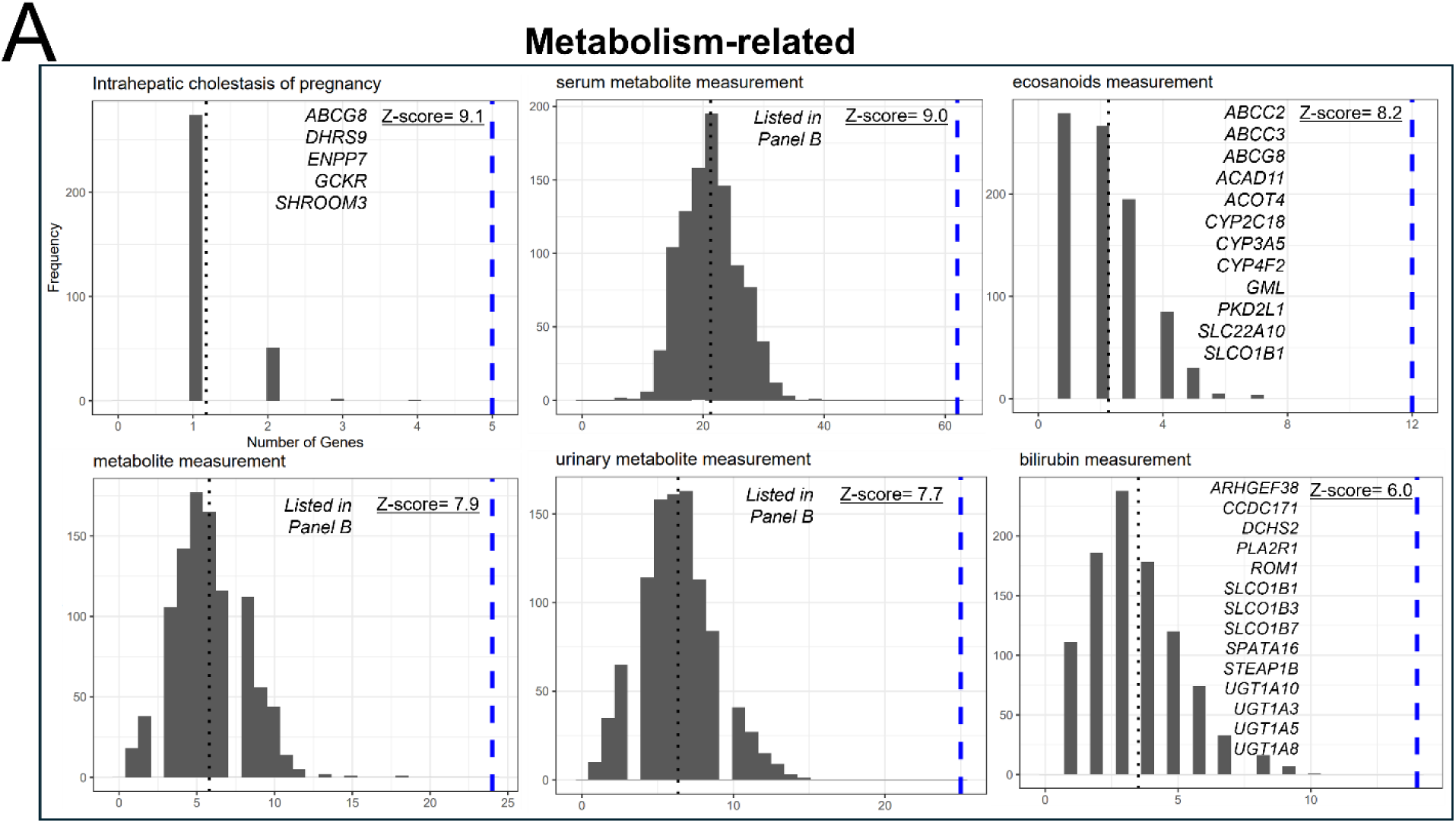
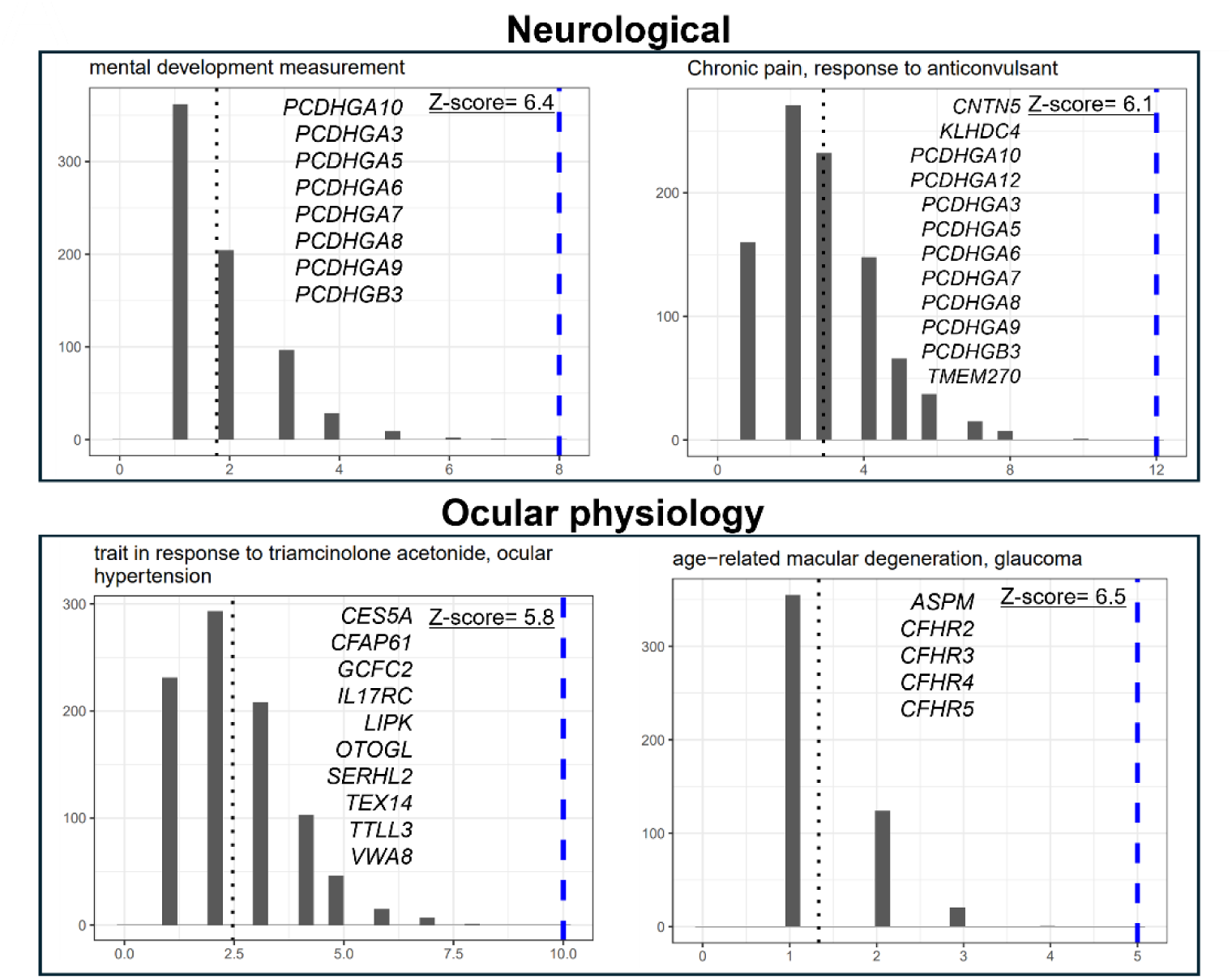
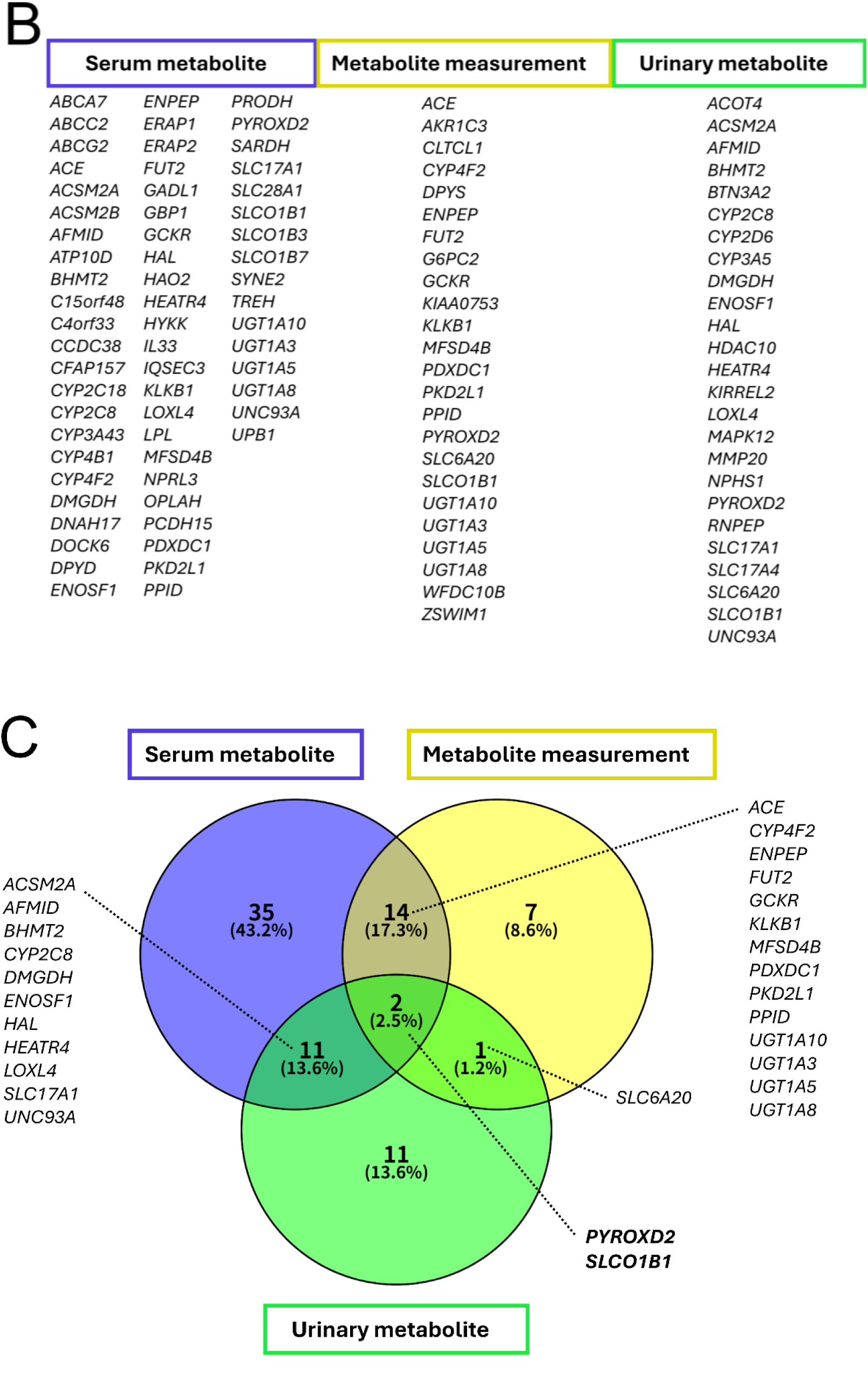
GWAS trait over-representation analysis of the PopVarLoF genes. The top 10 over-represented GWAS phenotypes ([GWAS Catalog](https://www.ebi.ac.uk/gwas/)) associated with the PopVarLoF genes were displayed in (A). In the individual histograms, the x-axis indicates the number of GWAS phenotype hits associated with the query gene sets (1551 PopVarLoF genes or 1551 random genes). The PopVarLoF genes are represented with the blue line while the average number of the random sets is represented with the black line. The y-axis indicates the corresponding frequency. The GWAS disease and trait categories are displayed at the top of each histogram plot. The overarching phenotype categories are also put on top of each GWAS phenotype (e.g., Metabolism-related, Ocular physiology, Neurological). Z-scores associated genes for GWAS phenotypes are also presented. Panel B lists genes associated with three GWAS phenotypes that include ‘metabolite’ in Panel (A). Panel (C) shows genes overlapping among the three GWAS phenotypes in Panel (B).

### 4. Over-representation of PheWAS-linked metabolic phenotypes

We similarly evaluated existing Phenome-wide association studies (PheWAS)(Ye et al., 2015b) in four databases (UKBTOPMED, KoGES, MGIBioVU, and FinMetSeq) for over-representation of phenotypic outcomes (Figure 5) associated with the PopVarLoF set. We identified 8 over-represented phenotypes associated with the PopVarLoF set in the UKBTOPMED data set, including “Poisoning by agents” (Z-score = 4.5), an aggregate trait of “Disorders of bilirubin excretion; Other disorders of metabolism; Other disorders of lipid metabolism; Lipoprotein disorders”, Z-score = 3.6, “Corns and callosities” (Z-score = 2.9), “Cervical cancer” (Z-score = 2.7), and “Disorder of skin and subcutaneous tissue” (Z-score = 2.6) (Figure 5A). The KoGES-based analysis identified 8 significant phenotypes, four of which were metabolism-related, including LDL Cholesterol (Z-score = 3.0), LDL Cholesterol (Meta-analysis) (Z-score = 2.5), Triglycerides (Meta-analysis) (Z-score = 2.4), and Total Cholesterol (Meta-analysis) (Z-score = 2.2) associated with the PopVarLoF genes. Other traits included Diastolic Blood Pressure (Z-score = 4.2), Diastolic Blood Pressure (Meta-analysis) (Z-score = 2.0), Hypertension (Z-score = 2.9), Hypertension (SPACox) (Z-score = 2.3) (Figure 5B). Both MGIBioVU and FinMetSeq databases-based analysis identified only one over-represented phenotype each and both were related to metabolism, Lactate dehydrogenase levels (in serum), (Z-score = 3.3) and Concentration of chylomicrons and extremely large VLDL particles (combined) (Z-score = 2.9), respectively. These results indicate over-representation of metabolism related phenotypes in the GWAS associations of the PopVarLoF set of genes.

**Figure 5.**
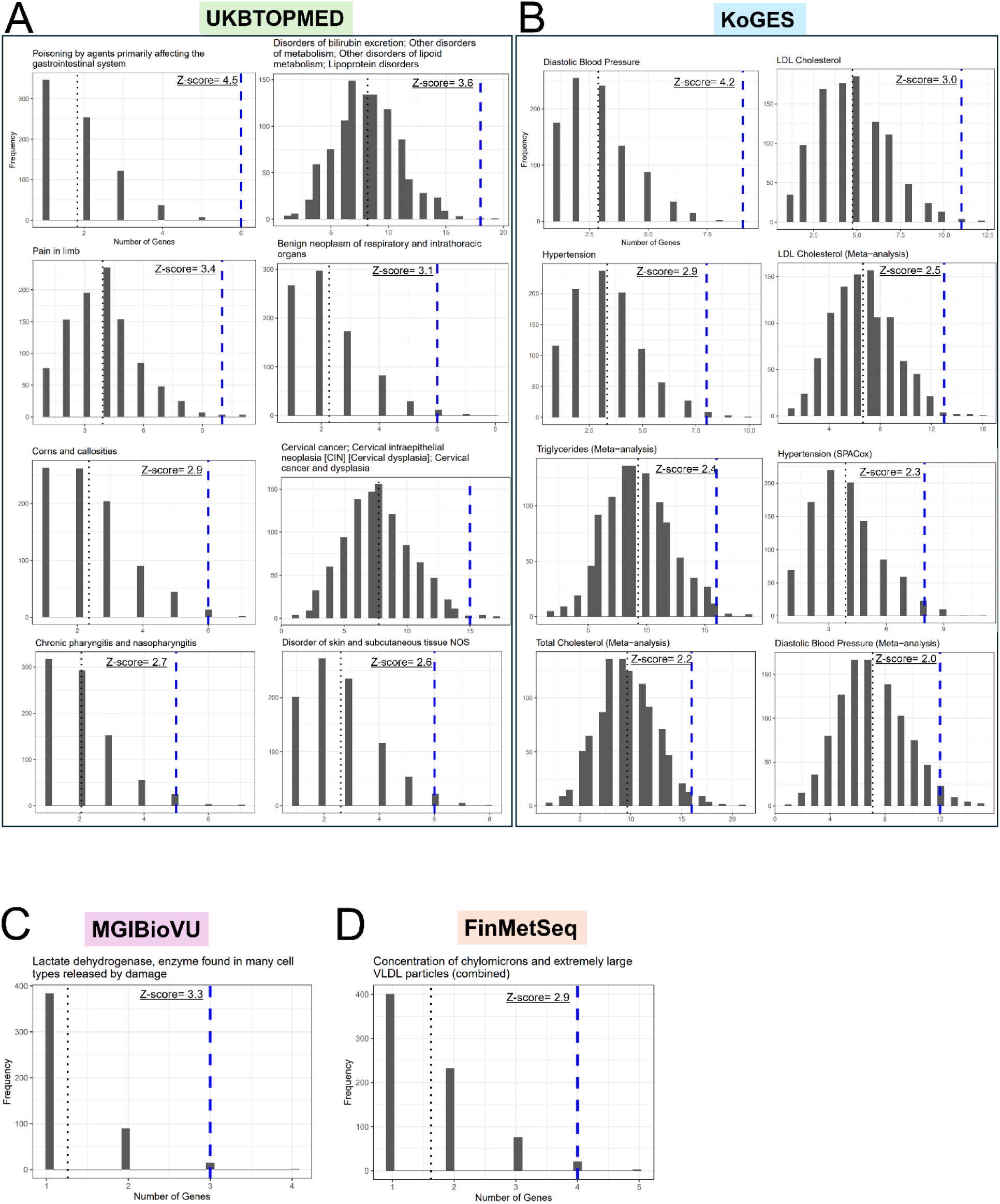
PheWAS trait over-representation analysis of the PopVarLoF genes. Significantly over-represented PheWAS phenotypes based on UKBTOPMED (A), KoGES (B), MGIBioVU (C), and FinMetSeq (D) are displayed. In the individual histograms, the x-axis indicates the number of PheWAS phenotype hits associated with the query gene sets (1551 PopVarLoF genes or 1551 random genes). The PopVarLoF genes are represented with the blue line while the average number of the random sets is represented with the black line. The y-axis indicates the corresponding frequency. The PheWAS disease and trait categories are displayed at the top of each histogram plot.

### 5. Functional requirement for PopVarLoF genes in HepG2/C3A

We assessed the effects of disruption of each gene in the PopVarLoF set on HepG2/CA3 proliferation and survival using a CRISPR KO screening approach (Figure 6A). We developed and validated custom PopVarLoF CRISPR KO libraries in LentiCRISPRv2 Puro, an all-in-one CRISPR vector with two different sgRNAs for each PopVarLoF gene (see methods). We carried out a CRISPR screen in the HepG2/C3A in the absence of any additional stressors other than two-dimensional cell culture. The HepG2/C3A cell line was transduced at low MOI, and we assessed depletion of each PopVarLoF mutant after approximately 7 cell doublings (T_7_) and 14 cell doublings (T_14_) of HepG2/C3A cells, respectively (Figure 6B) as compared to the initial cell population (T_0_). We identified significant depletion of 21 PopVarLoF mutants after T7 and 61 mutants after T_14_ with p-adj <0.05. We further identified a subset of 14 mutants (∼1% of the 1551 PopVarLoF set) which showed a trend of increased depletion from T_7_ to T_14_ (Figure 6B) consistent with their functional essentiality for continued survival in cell culture (Supplementary Figure S3 and Table S7). The 14 genes showed functional enrichment in mostly fundamental biological pathways (STRING analysis), including Catalytic activity acting on DNA (GO-BP), DNA helicase activity (GO-BP) and ribosomal RNA processing (STRING local), Ribosome biogenesis (STRING local) (Figure 6C).

**Figure 6.**
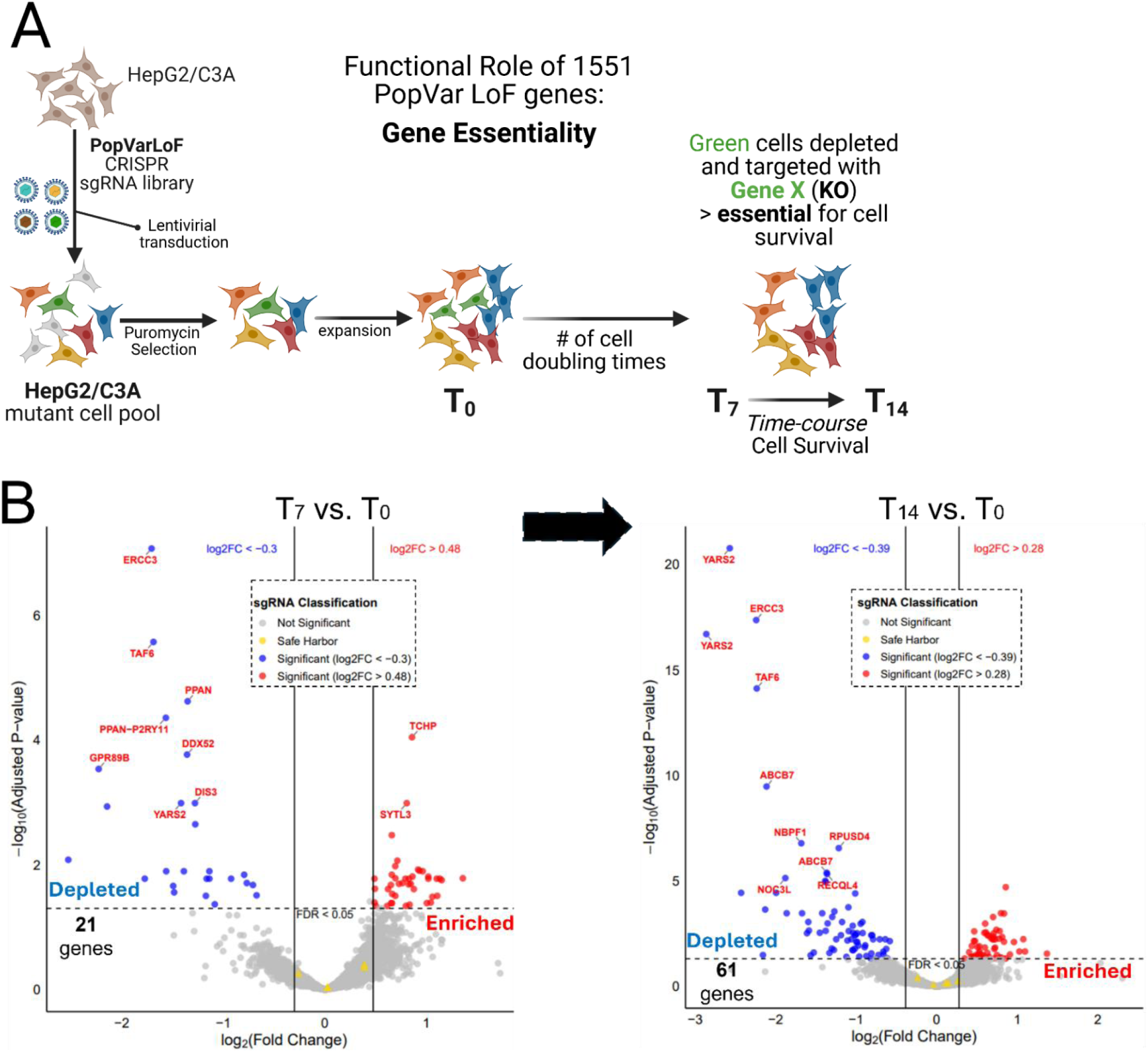
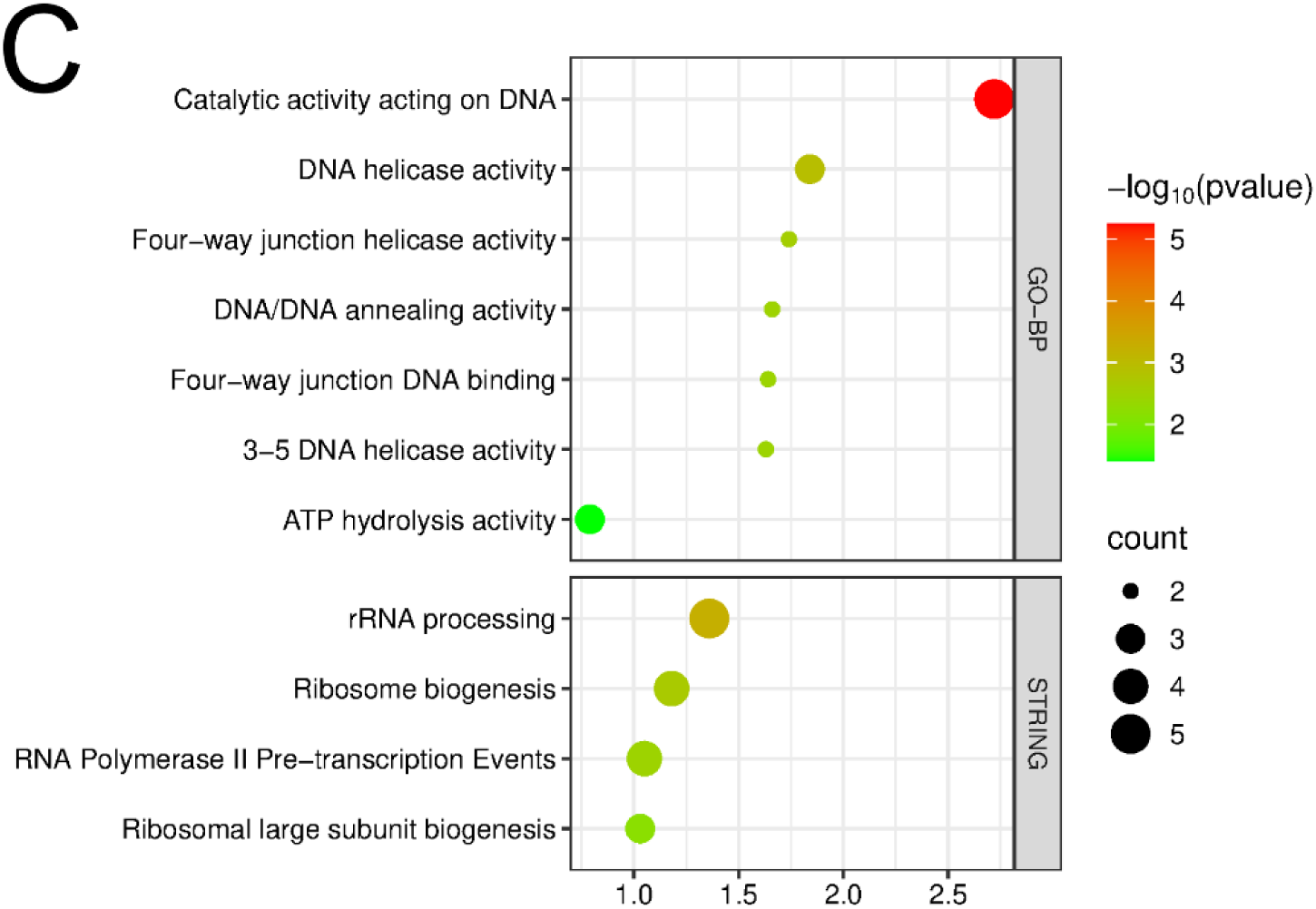
Essentiality of PopVarLoF genes in HepG2/C3A using CRISPR KO screening with PopVarLoF targeting library. (A) Experimental strategy to assess functional requirement of each PopVarLoF gene (referred to as Gene X) for normal growth and proliferation of HepG2/C3A by CRISPR KO screening. (B) Volcano plot of mutant KO pool comparing relative abundance of each KO mutant in pool initially (T_0_) and after an average of 7 doublings (T_7_) of the pool (Left panel). Similarly, a volcano plot compares relative abundance after 14 cell doublings (T_14_) to the T_0_ pool. (Right panel). A subset of genes showed increased depletion with Log_2_FC increasing from T_7_ to T_14_, Log_2_FC values < 0, and p-adj<0.05 for both Log_2_FC. (C) Dot plot for STRING enrichment result of the 14 essential PopVarLoF genes. The x-axis represents the enrichment signal, while the y-axis represents the enriched pathway term with pathway identification number (left to the plot) and the corresponding database (right to the plot). The color scale indicates different thresholds of the p-adj value, and the size of the dot indicates the number of genes corresponding to each pathway.

## Discussion

Variation in population and individual susceptibility to toxicants poses a considerable challenge to robust risk assessment to enable protection of susceptible individuals while simultaneously preventing biologically unrealistic predictions of adverse effects. Current risk assessment approaches to incorporate variability rely on the utilization of default uncertainty factors or probabilistic modeling(Varshavsky et al., 2023b). The regulatory and the scientific community have long recognized the need for more empiric measures of variability(National Research Council (US) Committee on Improving Risk Analysis Approaches Used by the U.S. EPA, 2009). In animal-based toxicity assays, the use of genetically diverse mouse strains, such as the diversity outbred (DO) set, can provide a surrogate estimate of inter-individual variability in toxicity effects in people(French et al., 2015; Saul et al., 2019). However, the lack of clear inter-species translations of observed variability, increased time and resources involved in toxicity testing in diverse rodent models make practical implementation in routine assessment challenging and unfeasible(Burnett et al., 2021; Rusyn et al., 2022). As toxicology and risk assessment transitions to the use of new approach methods (NAMs), including species appropriate *in vitro* cell-based systems, similar approaches based on toxicity testing in multiple cell lines derived from different individuals could enable sampling of genetic variability in NAMs based toxicity assays(Abdo et al., 2015a; Burnett et al., 2019; Ford et al., 2022). However, similar practical barriers to implementation include a requirement for testing of multiple individual cell lines with concomitant scaling of resource requirements, and a lack of clarity on the representation of an arbitrary set of individual cell lines of population level genetic variability, and uncertainty about the capture of biologically significant variants that could impact toxicity response(Abdo et al., 2015b; Dornbos and LaPres, 2018).

We propose a fundamentally different approach to this problem to estimate population relevant variation in toxicant susceptibly by systematically incorporating the most common predicted functional variants in the human population which could impact toxicity into human cell-based assays (Figure 7). We contend that a common functional variant or the set of common aggregate functional variants in a specific gene product, if they impact toxicant susceptibility of an individual, could have a significant population level effect on risk. While most identified genetic variants in the human population occur in non-coding, intergenic regions, and currently are primarily of uncertain functional significance(Burke et al., 2022), a subset of intragenic genetic variants are predicted to functionally disrupt the corresponding gene(MacArthur et al., 2012). Ongoing genome sequencing efforts compiled in the gnomAD database(Karczewski et al., 2020b; Gudmundsson et al., 2022) have systematically cataloged the frequency of both specific genetic variants predicted to impact the function of a gene but also the aggregate frequency of different genetic variants in a gene that could impair function(Karczewski et al., 2020b). In a risk assessment context, assessment of the most common functional variants in a human population could provide an empiric measure of the distribution of the genetically determined variance(Rusyn et al., 2022) in a cellular toxicity assay. We focus on the most common and most severe functional variants with the rationale that these variants, if impacting toxicity, could have a population level impact on risk. Specifically, we used a mean aggregate allele frequency >0.1% of a predicted loss of function genetic variants in a specific gene in a population as this MAF is used as the common individual variant cutoff in human genetic studies(Gudmundsson et al., 2022). We recognize that these common pLoF variants do not encompass all possible genetic variance impacting toxicity and additional less common variants of uncertain functional significance could be considered as technical capabilities make possible. The systematic introduction of population significant genetic variants provides an alternative, cost effective and scalable approach to assessing the impact of interindividual variability in any human cell-based assay system and if widely implemented could enable more data-driven estimates of population risk.

**Figure 7.**
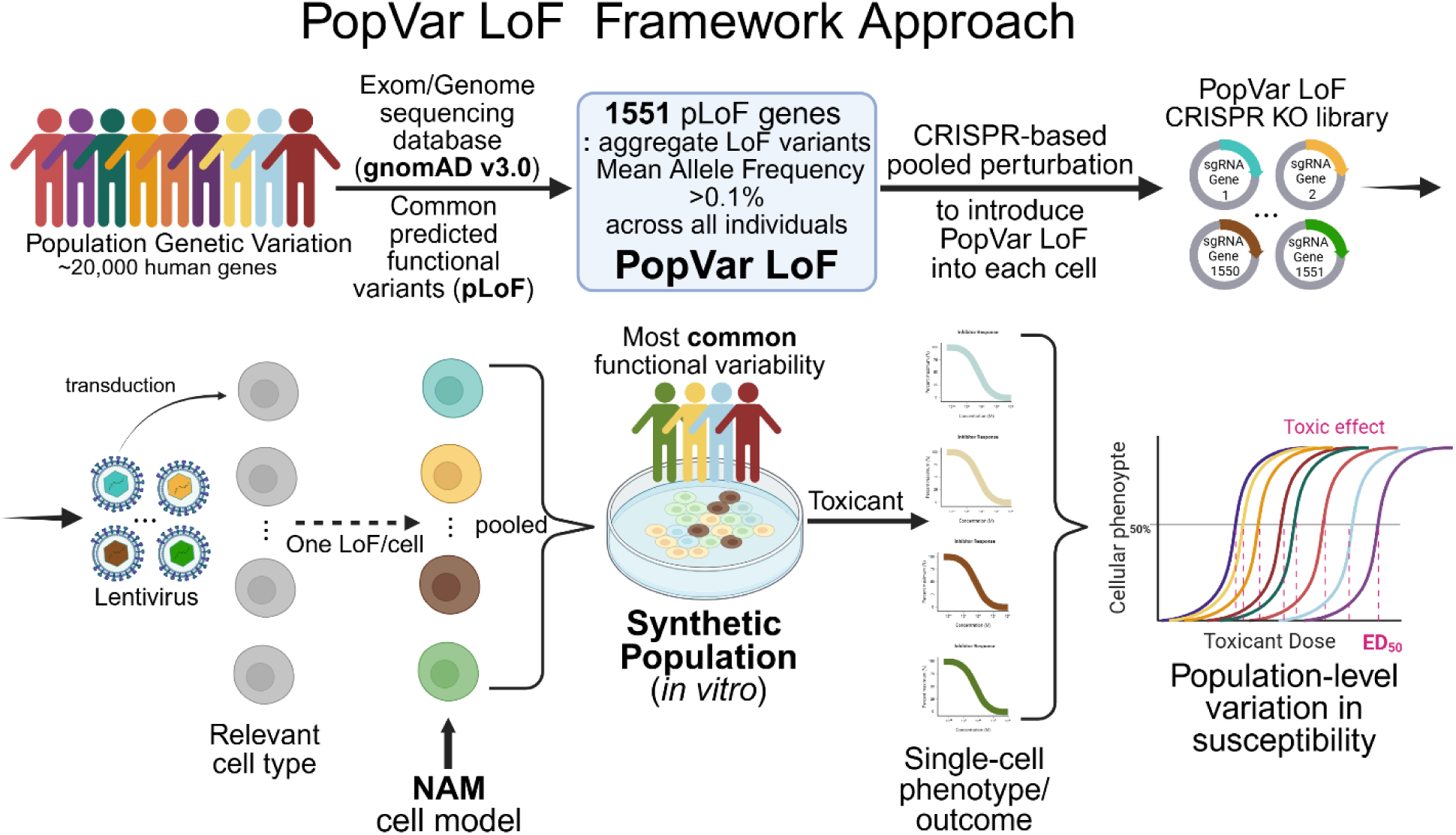
PopVarLoF framework for high-throughput experimental toxicological risk assessment proposed in this study. pLoF, predicted loss-of-function; KO, knockout; NAM, new approach methodology; ED_50_, effective dose that produces the desired effect in 50 % of the population being tested.

In this study, we identified and characterized the set of genes, which we labeled as the PopVarLoF set, in which pLoF variants met the criteria of mean aggregate allele frequency >0.1% as described above in the human population as of gnomAD v3.0. We recognize that the exact set of genes, and the predicted allele frequency will likely evolve as we gain additional genome sequence data and similarly, the population, for which the data is representative. Currently, genome sequence data is biased toward people of European descent(Sirugo et al., 2019) but ongoing efforts are attempting to provide more representative data sets(Auton et al., 2015; Fatumo et al., 2022). We initially focused on the set of pLoF which are common across the entire population set which may not be representative of the set of variants which are common in different sub-populations and thus may miss variants that are only common in one specific population. The development of population specific variation gene sets could enable targeted assessment of the distribution of toxicity susceptibility in different populations. Alternatively, weighted incorporation of common genetic variants from different population sub-sets into toxicity testing could enable more balanced representation.

Unexpectedly, the PopVarLoF set of genes was considerably enriched in gene products in xenobiotic, metabolism, and stress response related pathways previously reported for their roles in human toxicity responses(Vamvakas, 1997; Klaassen, 2002; Polonikov et al., 2009). Although speculative, LoF mutations in these, often induci(Lu et al., 2025)ble, gene products, may not impact reproductive fitness in the absence of environmental stressors(Rühli et al., 2016). However, this dispensability may come at a cost, as analysis of the entire gene set found considerable enrichment in both metabolism-related traits in GWAS associations and PheWAS phenotypes. Regardless of the cause, the representation of toxicant related pathways supports a plausible role, at least for represented genes, in underlying genetic variability in toxicant susceptibility, and suggests that functional interrogation of their role is warranted. As first step in this evaluation, we carried out an empiric assessment of their requirement for cellular essentiality(Wang et al., 2002) in one cell line model (i.e., HepG2/C3A) using CRISRP-based genetic screening system. We found that approximately 1 % (14 genes) of the PopVar genes were continuously depleted (measured by Log_2_FC: T_7_ → T_14_) in experimental cell populations in normal growth media while about 5.5 % of protein-coding genes covered in the DepMap(Tsherniak et al., 2017) database are defined to be essential. While the exact set of PopVar genes may evolve as additional data is obtained, and we learn more about their functional roles, the integration of these common pLoF variations into NAMs based cellular toxicity assays, particularly single cell-based phenotypes, will enable a more robust, scalable, approach to addressing the role of variability in genetic susceptibility in risk assessment.

Cost effective, high throughput and scalable incorporation of population variants into human cell-based assays will require development of novel approaches.(Chu et al., 2016; Jovic et al., 2022; Stossi et al., 2023). We adapted CRISPR KO based screening approach(Shalem et al., 2014; Wang et al., 2014) to individually disrupt each gene with a pLoF aggregate MAF of >0.1% in the human population of interest in the assay cell line (HepG2/C3A in this study) in such a way that a single gene is targeted in each cell but collectively all potential pLoF variants are represented in a population of cells(Doench, 2018). Thus, a “synthetic population” of cells which phenocopies the predicted functional effect of a homozygous LoF in each gene. Importantly, each cell contains at most a single LoF in a single gene in a common genetic background so any observed change in individual cellular phenotype can be attributed to the introduced variation. A synthetic cellular population with different individual variants in combination with single cell-based phenotypes could enable simultaneous assessment of point of departure (POD) of each LoF variant on each cellular toxicity endpoint. Similarly, the distribution POD could enable estimation of the toxicodynamic variability factor at 5% (TDVF_05_) across the synthetic population representative of population relevant variants(Blanchette et al., 2022). If the goal is to estimate the distribution of effects of common pLoF variants on one or more toxicity endpoints, the identity of the introduced LoF variation in each cell in the synthetic population may not be necessary and high throughput single cell phenotype assessment can be utilized without knowing the identity of each pLoF variant in each cell. If the goal is to link a specific LoF variation with a specific toxicity outcome, then identification of each variant would be required. While not currently routine, evolving optical CRISPR technology enables linking specific variant introduction and phenotypic outcome currently through various approaches including *in situ* sequencing(Kahnwald et al., 2024). Alternatively, if the phenotypic outcome is RNA based, such as transcriptional induction, coupled CRISPR-single cell RNA-seq based approaches allow attribution of RNA changes to specific introduced genetic changes(Dixit et al., 2016; Cheng et al., 2022; Replogle et al., 2022). We recognize that isolation of cell lines containing specific variants and subsequent individual testing of each variant cell line is possible(Kim and Vulpe, 2026) but would require the parallel testing of multiple variant cell lines, each with a different variant, and thus lead to similar issues with scalability as testing multiple cell lines from different individuals. We anticipate that targeted subsets may be possible as the toxicology community identifies the common variants most likely to impact toxicity phenotypes.

## Supplementary material

Supplementary material is available at *Toxicological Sciences* online.

## Data availability

The raw and processed CRISPR screen data with the corresponding metadata in this study will be deposited into Gene Expression Omnibus (GEO) database repository as soon as GEO submission is available.

