## Supplementary Figures for "Characterizing common loss-of-function genes and their potential utility in assessing population variability and chemical susceptibility"

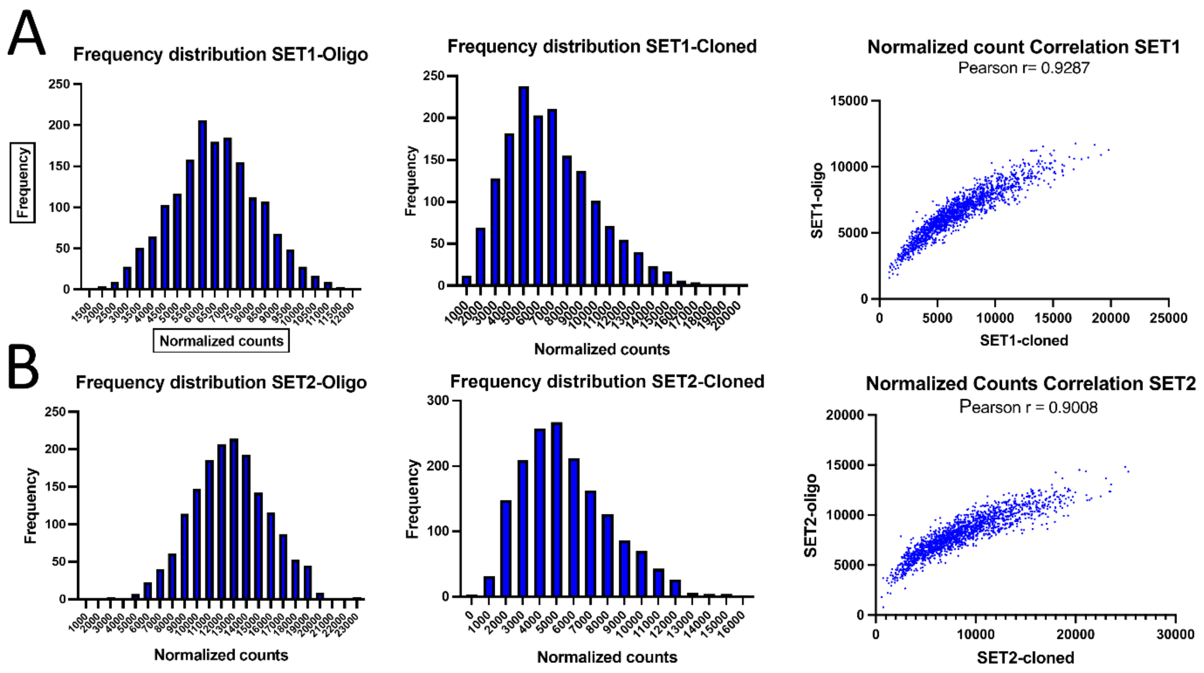


**Figure S1**. Quality assessment of the custom sgRNA libraries (Set1 and Set2) was conducted. The distribution of normalized sgRNA counts and their corresponding frequencies in the library before and after

cloning (Oligo-pool vs. cloned) are shown in panels A and B. The correlation between the Oligo-pool and the cloned library is presented with the Pearson correlation coefficient (r) in panel C.

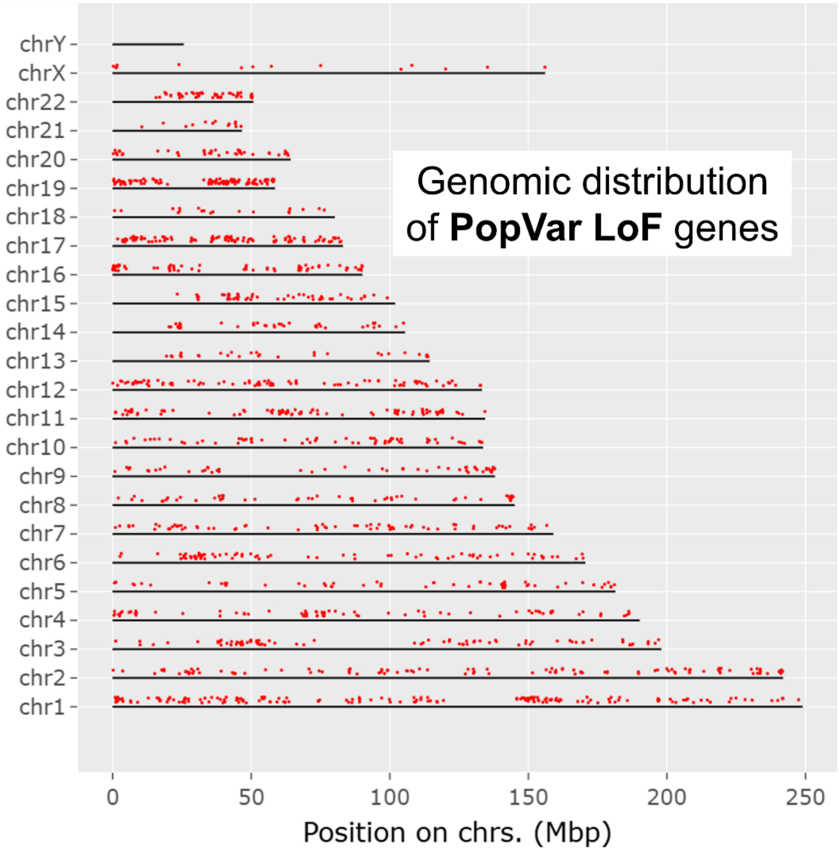


**Figure S2.** Distribution of individual PopVarLoF genes across chromosomes.

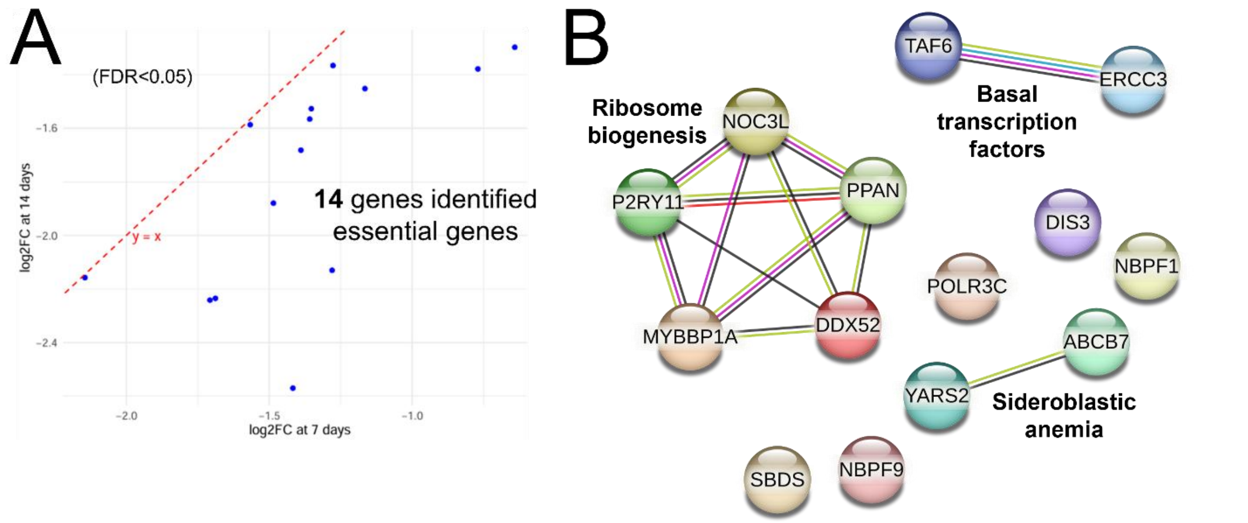


**Figure S3**. Empirically identified essential genes (14 genes) in the PopVar LoF CRISPR screening in a time-course manner (7 doublings: $T_7$ and 14 doublings: $T_{14}$). (A) Blue dots indicate 14 essential PopVarLoF genes which showed increased depletion between $T_7$ and $T_{14.}$ (B) STRING functional enrichment of the 14 essential genes. Three clusters were identified as presented Ribosome biogenesis, Basal transcription factors, and Sideroblastic anemia. Detailed features on these 14 genes are available in Table S3.
